# Deubiquitinase UCHL1 Maintains Protein Homeostasis through PSMA7-APEH-Proteasome Axis in High-Grade Serous Ovarian Carcinoma

**DOI:** 10.1101/2020.09.28.316810

**Authors:** Apoorva Tangri, Kinzie Lighty, Jagadish Loganathan, Fahmi Mesmar, Ram Podicheti, Chi Zhang, Marcin Iwanicki, Harikrishna Nakshatri, Sumegha Mitra

## Abstract

High-grade serous ovarian cancer (HGSOC) is characterized by chromosomal instability, DNA damage, oxidative stress, and high metabolic demand, which exacerbate misfolded, unfolded and damaged protein burden resulting in increased proteotoxicity. However, the underlying mechanisms that maintain protein homeostasis to promote HGSOC growth remain poorly understood. In this study, we report that the neuronal deubiquitinating enzyme, ubiquitin carboxyl-terminal hydrolase L1 (UCHL1) is overexpressed in HGSOC and maintains protein homeostasis. UCHL1 expression was markedly increased in HGSOC patient tumors and serous tubal intraepithelial carcinoma (HGSOC precursor lesions). High UCHL1 levels correlated with higher tumor grade and poor patient survival. UCHL1 inhibition reduced HGSOC cell proliferation and invasion through the outer layers of omentum as well as significantly decreased the *in vivo* metastatic tumor growth in ovarian cancer xenografts. Transcriptional profiling of UCHL1 silenced HGSOC cells revealed the down-regulation of genes implicated with proteasome activity along with the upregulation of endoplasmic reticulum (ER) stress-induced genes. Reduced expression of proteasome subunit alpha 7 (PSMA7) and acylaminoacyl peptide hydrolase (APEH) resulted in a significant decrease in proteasome activity, impaired protein degradation, and abrogated HGSOC growth. Furthermore, the accumulation of polyubiquitinated proteins in the UCHL1 silenced cells led to attenuation of mTORC1 activity and protein synthesis, and induction of terminal unfolded protein response. Collectively, these results indicate that UCHL1 promotes HGSOC growth by mediating protein homeostasis through the PSMA7-APEH-proteasome axis.

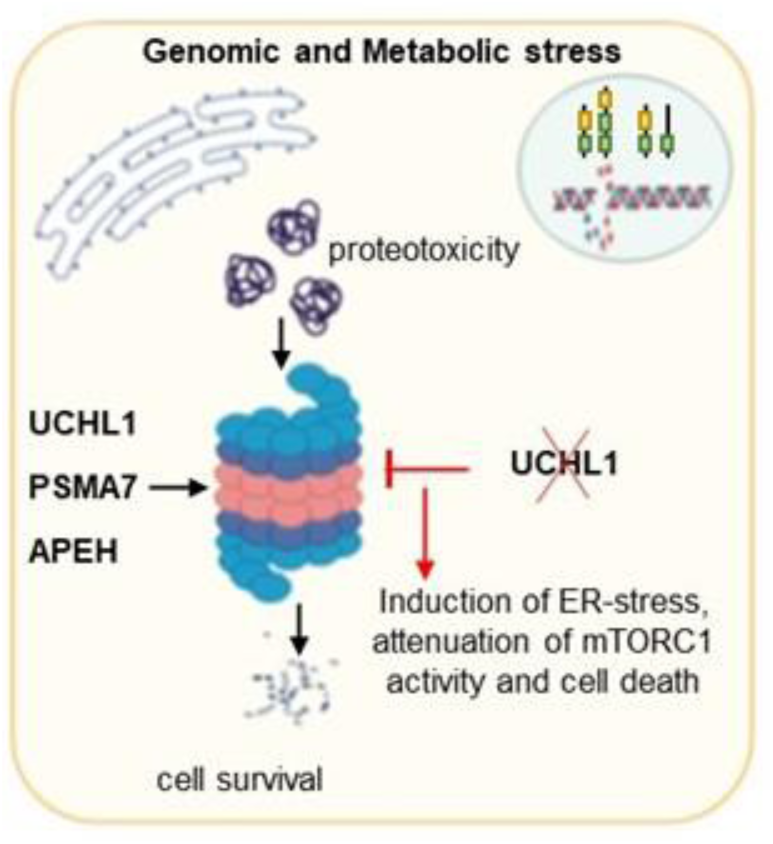

**Implications:** This study identifies the novel links in the proteostasis network to target protein homeostasis in HGSOC. It recognizes the potential of inhibiting UCHL1 and APEH to sensitize cancer cells to proteotoxic stress and as novel alternative therapeutic approaches.

## Introduction

Cancer cells maintain protein homeostasis to sustain their high proliferating state. Protein synthesis is intrinsically an error-prone process and up to 30% of newly synthesized misfolded proteins are degraded immediately after protein translation [1, 2]. Moreover, Cancer cells with profound chromosomal instability, mutations, and physiological stressors carry the burden of excessive protein production, mutant proteins with stoichiometrically altered protein complexes, and increased misfolded and damaged proteins [3-5]. Together, this contributes to a proteotoxic state in cancer cells, if misfolded or damaged proteins are not efficiently removed [2, 4-6]. Therefore, understanding the mechanisms that regulate protein homeostasis is an essential link to develop effective treatment strategies. Disrupting this equilibrium through the use of proteasome inhibitors has already revolutionized the treatment of hematological malignancies, such as multiple myeloma and mantle cell lymphoma [7]. However, the first-generation proteasome inhibitor, Bortezomib has shown limited success in solid tumors [7], suggesting the need for alternative approaches to specifically target protein homeostasis in solid tumors.

The ubiquitin-proteasome system is at the core of the protein quality control network and works together with protein folding and protein clearance pathways to maintain protein homeostasis [2, 5]. Most cancer cells display enhanced proteasome activity to maintain the integrity of the onco-proteome, regulate cellular levels of proteins like cell-cycle checkpoints or tumor suppressors, and avoid growth arrest due to the accumulation of misfolded proteins [7, 8]. Proteasome inhibition induces an integrated stress response as a result of amino acids and ubiquitin deprivation, reduced protein synthesis, and increased endoplasmic reticulum (ER) stress, which induces terminal unfolded protein response [9-11]. It is now clear that cancer cells adapt in various ways to maintain protein homeostasis and enhance proteasome activity through upregulation of proteasome subunits, proteasome activators, or proteasome assembly factors [12-15], which makes them fascinating selective targets to block proteasome activity in cancer cells. Emerging in this field are deubiquitinating enzymes (DUBs) inhibitors [16]. A pan-deubiquitinating enzyme inhibitor has been reported to sensitize breast cancer cells to the proteotoxicity caused by oxidative stress in absence of glutathione [16]. Moreover, the small-molecule inhibitor of proteasome-associated DUBs, b-AP15 has been reported to overcome Bortezomib resistance, inducing proteotoxic stress, and reactive oxygen species (ROS) [17]. These studies demonstrated the effect of global DUB inhibition on proteotoxic stress-induced cancer cell death, however, the knowledge of a specific DUB remains elusive.

Ubiquitin carboxyl-terminal hydrolase L1 (UCHL1) is a neuronal DUB, that constitutes about 1-2% of total brain proteins [18]. The loss of UCHL1 has been implicated in the accumulation of neuronal protein aggregates due to impaired proteasomal degradation in neurodegenerative diseases [19, 20]. Though UCHL1 is overexpressed in several malignancies [21-23], nothing is known about its role in the protein clearance pathway in cancer. UCHL1 plays a promiscuous role in cancer and has been shown to promote metastatic growth by its deubiquitinating activity associated with HIF1α, cyclin B1, and TGFβ receptor 1 [21, 22, 24], while it is reported as an epigenetically silenced tumor suppressor in some cancers [25, 26]. In the present study, we report that increased expression of UCHL1 in high-grade serous ovarian cancer (HGSOC) mediates protein homeostasis. HGSOC is the most prevalent and lethal histotype of ovarian cancer. It is characterized by chromosomal instability, germline, or somatic mutations, including the mutations in the tumor suppressor gene *TP53* [27]. However, not much is known about the mechanisms that mediate proteostasis in HGSOC. Here we show that UCHL1 mediates protein homeostasis through increased acylaminoacyl peptide hydrolase (APEH) activity and proteasome subunit alpha 7 (PSMA7) expression. Furthermore, UCHL1 inhibition results in the accumulation of polyubiquitinated proteins leading to the induction of terminal unfolded protein response and attenuation of mTORC1 (mTOR complex 1) activity and protein synthesis. This is the first report to establish the role of UCHL1 in mediating protein homeostasis through the PSMA7-APEH-proteasome axis and identifies the novel druggable links to target protein homeostasis in HGSOC.

## Materials and Methods

### Cell culture

All ovarian cancer cell lines were maintained in 10% DMEM medium (Corning; cat#10-013-CV) supplemented with 1% non-essential amino acid and 1% vitamins and 1% penicillin/streptomycin (Corning). Fallopian tube (FT) epithelial cells were obtained from Dr. Ronny Drapkin, University of Pennsylvania, and were cultured in DMEM-Ham’s F12 media (Corning) supplemented with 2% UltroserG (Crescent Chemical Company). Non-ciliated FT epithelial (FNE) cells transfected with vector pWZL-mutant p53-R175H were maintained in WIT-Fo Culture Media from Live Tissue Culture Service Center, University of Miami by the laboratory of Dr. Marcin Iwanicki. Human primary mesothelial cells (HPMC) and fibroblasts isolated from the omentum of a healthy woman were obtained from Dr. Anirban Mitra, Indiana University, and were grown in 10% DMEM. All cell lines were authenticated by short tandem repeat (STR) profiling and were negative for mycoplasma contamination.

### Patient samples and patient data analysis

Frozen human serous ovarian cancer primary tumors and matched normal adjacent fallopian tubes (FT) were obtained from the tissue bank of Indiana University Simon Cancer Center (IUSCC). The study was approved by the Institutional Regulatory Board of Indiana University (protocol numbers 1106005767 and 1606070934). Human serous tubal intraepithelial carcinomas tissue slides (n=3) were obtained from Dr. Marcin Iwanicki, Stevens Institute of Technology. Tissue microarrays of HGSOC tumors with normal ovary (OV1502 and BC11012) and normal fallopian tube (UTE601) were purchased from US Biomax Inc and were processed at the same time. Written consent was obtained from all the patients and only de-identified patient specimens were used. TCGA database was analyzed using the Oncomine gene browser [28] to examine gene expression in HGSOC patients and across cancer stages and tumor grades. Survival analysis of HGSOC patients (n=1104) who had received chemotherapy after optimal or suboptimal debulking was performed using the KM plotter [29]. Survival analysis of HGSOC patients was analyzed in an *in-house* cohort of Molecular Therapeutics for Cancer, Ireland (MTCI), and GSE9899 (n=244) using OVMARK [30]. Patients with no residual tumors and UCHL1 median expression were used as the cut-off. Correlation between UCHL1 and p*53* expression levels in HGSOC patients with *TP53* mutations (putative driver n=92, missense mutation n=143, and no mutation n=10) was analyzed in TCGA database using cBioportal [31].

### Animal study

The animal study was performed according to protocols approved by the Animal Care and Use Committee of Indiana University. Five million OVCAR8 cells were intraperitoneally injected into 5-6-week old female athymic nude mice (Envigo) as described earlier [32]. Mice were randomized into two groups: vehicle control and LDN5777 (10 mice/group). After 10 days of injecting the cancer cells, mice were intraperitoneally injected with LDN57444 (1mg/Kg) or 25% DMSO thrice/week for 5 weeks. All the mice were euthanized after 45 days of injecting the cells.

### Methylated DNA immunoprecipitation (MeDIP)

MeDIP was performed using the Active Motif kit (Cat# 55009). The genomic DNA was isolated from the ovarian cancer cell lines using the DNeasy Blood and Tissue kit (Qiagen). DNA (20ng/µl) was sheared on ice for 3 pulses of 10 seconds at 30% amplitude with a 20 second pause between each pulse using a tip probe sonicator. DNA fragment size was ensured by Agilent TapeStation. MeDIP was performed using 5-methylcytosine antibody or control mouse IgG according to the manufacturer’s protocol. Quantitative PCR was performed in the input and MeDIP samples for UCHL1 promoter using primers, forward: ccgctagctgtttttcgtct and reverse: ctcacctcggggttgatct. The analysis was performed as a percent of input normalized to control IgG. Amplicons were resolved using a 2% agarose gel.

### Chromatin immunoprecipitation (ChIP)

ChIP was carried out using the ChIP-IT Express kit (Active Motif, cat# 53008). Briefly, cells were fixed in 1% methanol-free formaldehyde (Fisher Scientific, cat# 28908) followed by Glycine-Stop Fix solution treatment. Cells were lysed as per the manufacturer’s protocol. The nuclei were suspended in the shearing buffer and sonicated for 8 cycles of 30sec on/off using Bioruptor Pico (Diagenode). The sheared chromatin was reverse-crosslinked and DNA fragment size was ensured by Agilent TapeStation. ChIP was performed according to the manufacturer protocol using an anti-histone H3K4 trimethyl antibody (Abcam, ab8580) or control IgG. Quantitative PCR was performed for UCHL1 promoter using primers ccgctagctgtttttcgtct, ctcacctcggggttgatct. The analysis was performed using the 2^-ΔΔCt^ method [33].

### Cell proliferation and colony formation assay

Cell proliferation was measured by MTT assay as described earlier. 2000 cells transfected with control or target specific siRNA per well were plated in the 96-well plate and MTT assay was performed after on day 4. The reduction of MTT into purple color formazan was measured at 560nm and adjusted for background absorbance at 670nm. Colony formation assay was performed by plating 1000 cells per well in the 6-well plate. The colonies were allowed to grow for 8-10 days and the fixed colonies were stained with 0.05% crystal violet solution. The colonies were imaged and counted using ImageJ.

### Spheroid culture of FNE cells and LDN57444 treatment

Fallopian tube non-ciliated epithelial (FNE) cells transfected with pWZL-p53-R175H to overexpress mutant p53 variant R175H (FNE^mutp53-R175H^) and green-fluorescence protein (GFP) were seeded in ultra low-adhesion plates (Corning). 2% Matrigel was added to the suspended culture after 24 hours to support basement membrane adhesion. After 4 days, the three-dimensional (3D) structures of FNE^mutp53-R175H^ cells were treated with DMSO or UCHL1 inhibitor, LDN57444 (10 µM, 5 days). Subsequently, cellular clusters were treated with 2μM ethidium bromide (EtBr) and were imaged. EtBr incorporation was measured as the number of red channel pixels within cellular clusters as described earlier [34].

### Organotypic three-dimensional (3D) culture model of omentum and invasion assay

The organotypic 3D culture model of the omentum was assembled in a fluoroblock transwell insert (8µm pore size, BD Falcon) as described earlier [35]. Briefly, 2×10^5^ fibroblasts with collagen I and 2×10^6^ primary mesothelial cells isolated from the omentum of a healthy woman were seeded in the transwell insert. After 24 hours, 2×10^5^ UCHL1 silenced or unsilenced OVCAR3 (RFP labeled) and Kuramochi (GFP labeled) cells were plated over the omental cells in 200µl of serum-free DMEM. Cancer cells were allowed to invade for 16 hours after placing the insert in a well of 24-well plate containing 700μl of 10% DMEM. Invaded cells were fixed, imaged (5 fields/insert), and counted.

### Determination of proteasome and APEH activity

The chymotrypsin-like proteasome activity was measured by Sigma-Aldrich kit (cat# MAK172) using the fluorogenic substrate LLVY-R110 as per the manufacturer’s protocol and as described earlier [36]. Total protein (50µg) from fresh cell lysate or tissue homogenates in even volumes (90 µl) was incubated with 100µl of proteasome assay buffer containing LLVY-R110 at 37°C. R110 cleavage by proteasomes was measured at 525nm with excitation at 490nm. Fluorescence intensity was normalized with the fluorescence of blank well. APEH activity was measured by chromogenic substrate acetyl-Ala-pNa (Bachem) [37]. Total protein (45µg) in even volume (100 µl) in 50mM Tris-HCl buffer pH7.5 was incubated with acetyl-Ala-pNa at 37°C. The release of p-nitroaniline was measured at 410nm and was normalized with the absorbance of the blank well.

### Immunoblot analysis

Immunoblotting was performed using a standard protocol as described earlier [35]. Cells were lysed in NP-40 buffer containing protease and phosphatase inhibitors cocktails (Millipore), 0.2mM phenylmethylsulfonyl fluoride, and 10mM N’ ethylmalamide. Protein quantification was conducted using the Pierce BCA protein assay kit (Thermo Fisher #23225). Proteins were resolved by 4-20% gradient SDS-PAGE. Primary antibodies used were UCHL1 (13179, Cell Signaling), UCHL1 (MAB6007, R&D Systems), PSMA7 (cat# PA5-22289; Invitrogen), ATF3 (cat# 33593; Cell signaling), APEH (cat# 376612; Santacruz Biotechnology), and actin-HRP (Sigma).

### Transfection, transduction, and cell treatments

Gene knockout was carried out by transfecting HGSOC cells with control and target specific siRNAs (set of four siRNAs) Dharmacon ON-TARGETplus siRNA (Horizon Discovery) for UCHL1 (cat# L-004309-00-0010), PSAM7 (cat# L-004209-00-0010), APEH (cat# L-005785-00-0010) and control (cat# D-001810-10-05) using TransITX2 (Mirus Bio; cat#MIR6000). UCHL1 knockdown was also carried out by transducing HGSOC cells with control or UCHL1 shRNA lentiviral virus particles (Santacruz Biotechnology; cat# sc-108080 and sc-42304-V) using TransDux™ MAX (System Biosciences; cat#LV860A-1). Cells were treated with Carfilzomib (cat# S2853, Selleck Chemicals) and 5-Aza-2′-deoxycytidine (Sigma-Aldrich; cat# A3656) at the indicated dose and time points. For 5-Aza-2′-deoxycytidine (5µM; 48h) [38] treatment cells were plated at a low density that will allow its incorporation into the DNA of the dividing cells.

### Immunohistochemistry (IHC)

IHC was performed by IU Health Pathology Laboratory. Briefly, slides were baked at 60°C for 30 minutes before the standard deparaffinization procedure followed by blocking of endogenous peroxides and biotin. Antigen retrieval was performed using 10mM citrate buffer, pH6.0 at 95°C followed by 1h blocking and incubation with pre-optimized primary anti-UCHL1 (MAB6007, R&D Systems) or anti-p53 (Dako) antibodies (1:200 dilution). TMA slides were digitally scanned by Aperio ScanScope CS slide scanner (Aperio Technologies) and staining was quantified in three intensities ranges: weak – 0 to 100, positive – 100 to 175, and strong – 175 to 220. TMA slides were also hand-scored by Dr. George Sandusky as 1 being a weak expression, 2 moderate, 3 strong, and 3+ very strong.

### Assay for transposase-accessible chromatin (ATAC) sequencing

ATAC-seq was performed by the Center for Medical Genomics, Indiana University School of Medicine. The Tagment DNA TDE1 enzyme and Nextera DNA Flex Library Prep kit (Illumina cat#15027866 and 15027865) were used. Briefly, 1×10^5^ OVCAR3 and SKOV3 cells were lysed in a non-ionic detergent to yield pure nuclei. The chromatin was fragmented and simultaneously tagmented with the sequencing adaptor using Tn5 transposase to generate ATAC-seq libraries, which were sequenced on NextSeq 500 (Illumina) with NextSeq75 High Output v2 kit (Illumina, cat# FC-404-2005). Raw fastq files were aligned to the human GRCH38 genome by using bowtie 2 [39]. MACS2 and ENCODE standardized pipeline and parameters were utilized for peak detection [40]. Peaks on the promoter region of the UCHL1 gene were plotted using the UCSC genome browser [41].

### RNA isolation, real-time PCR, and RNA sequencing

Total RNA from cell lines and patient tumors was extracted using the miRNeasy mini kit (Qiagen; cat#217004). Real-time PCR was performed using TaqMan gene expression assays after cDNA preparation using the high capacity reverse transcription kit (Applied Biosystems; cat#4368814). β-actin and tata-box binding protein (TBP) were used as endogenous controls. For RNA-sequencing, 1 µg of total RNA was used for library preparation using the TruSeq Stranded mRNA kit (Illumina, cat# RS-122-2103) after rRNA depletion using Ribo-Zero plus (Illumina; cat#20037135). RNA-sequencing was performed using NextSeq75 High Output v2 kit and NextSeq 500 (Illumina; cat# FC-404-2005). Using TruSeq 3’ SE adaptor sequence AGATCGGAAGAGCACACGTCTGAACTCCAGTCAC, RNA-seq reads were trimmed and then were mapped to the gene regions in a strand-specific manner using htseq-count (version 0.5.4p1) [42]. Differentially expressed genes at 5% FDR with at least two-fold change were called using DESeq2 ver.1.12.3 as described earlier [43].

### Statistical analysis

Statistical significance was calculated using Student *t*-test and one-way ANOVA using Prism 8.0. The logrank test was used to determine the significance of survival analysis. All results are expressed as mean ± SD from three biological repeats unless otherwise stated. The *p-*value of less than 0.05 was considered significant.

## Results

### UCHL1 overexpression is an early event in HGSOC and associates with poor patient prognosis

To assess the role of UCHL1 in HGSOC, we examined publicly available TCGA data of serous ovarian cancer patients. Our analysis of TCGA data revealed that UCHL1 is a frequently overexpressed gene in HGSOC patients. UCHL1 mRNA levels were significantly high in primary and recurrent tumors compared to normal ovaries (Figure 1A). Moreover, UCHL1 expression was markedly elevated in HGSOC patients with advanced-stage and higher-grade tumors compared to grade 1 and stage 1 tumors, respectively (Figures 1B and 1C). To confirm these results at the protein level, we performed UCHL1 immunohistochemical (IHC) staining in tissue microarrays consisting of HGSOC tumors, normal ovaries and normal fallopian tubes (FT). Compared to normal tissues, UCHL1 expression was significantly higher in HGSOC tumors (Figures 1D and 1E). UCHL1 staining was positive in 78.4% (69 out of 88) of tumors and its expression was high in 59.1% (52 out of 88) of tumors, whereas its expression was negligible or low in the normal fallopian tube (FT) and ovary. Furthermore, UCHL1 mRNA and protein levels were elevated in primary tumors compared to their matched normal adjacent FT (Figure 1F, paired samples). These results suggest that UCHL1 expression is upregulated in HGSOC. To test if UCHL1 expression is an early event in HGSOC, we performed UCHL1 IHC staining in serous tubal intraepithelial carcinoma (STIC). HGSOC is known to originate from the lesions in FT known as STICs. *TP53* mutations are an early event in the development of STICs and the presence of identical *TP53* mutations in STICs and concurrent HGSOC established their clonal relationship [44]. UCHL1 levels were significantly elevated in the STICs as evidenced by the increased UCHL1 staining in the epithelial cells and the associated invasive carcinoma with diffused nuclear staining of mutant p53 (Figure 1G), while the UCHL1 staining was absent in p53-negative regions and normal human FT (Figures 1G and S1). Next, to determine the prognostic significance of UCHL1, we analyzed the transcriptomic datasets of HGSOC patients. Survival analysis using the Kaplan Meier plotter revealed a significant association of high UCHL1 levels with poor progression-free survival of HGSOC patients after chemotherapy and debulking (Figure 1H). Moreover, high UCHL1 levels correlated with poor disease-free survival of HGSOC patients after optimal debulking in the survival analysis of GSE9899 and an *in-house* cohort of Molecular Therapeutics for Cancer, Ireland using OVMARK (Figure 1I). Overall, these results indicate that UCHL1 overexpression in HGSOC patient tumors is an early event and predicts poor prognosis, indicating its essential role of in HGSOC.

**Figure 1.**
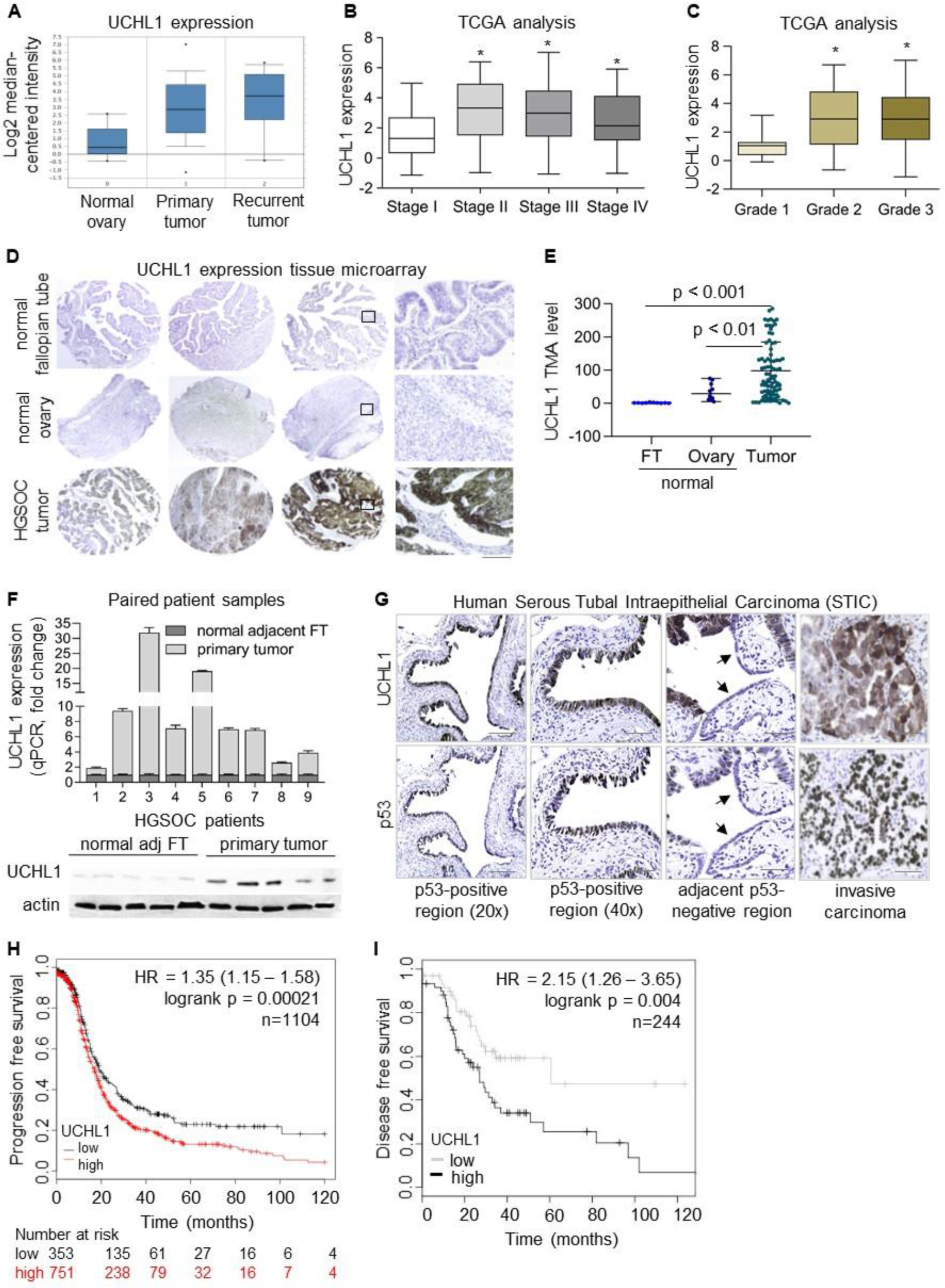
UCHL1 overexpression confers poor prognosis in HGSOC patients. **A**. UCHL1 mRNA expression in primary and recurrent tumors of HGSOC patients in TCGA database analyzed using the Oncomine gene browser. **B**. UCHL1 expression in stage I (n=16), stage II (n=27), stage III (n=436), and stage IV (n=84) tumors of HGSOC patients in TCGA database. **C**. UCHL1 expression in grade 1 (n=15), grade 2 (n=69) and grade 3 (n=479) tumors of HGSOC patients in TCGA database. **D**. Representative core images for low, medium, and high UCHL1 levels in HGSOC tumors (n=88), normal fallopian tube (FT), and normal ovary (n=10 each) in the tissue microarray (TMA) of HGSOC patients, scale bar: 200µm and 50µm. **E**. Quantification of UCHL1 expression (H-score) by digital scanning of TMA. **F**. Relative UCHL1 mRNA and protein levels in primary HGSOC tumors and matched normal adjacent fallopian tubes obtained from the same patient (n=9 pairs). Top: qPCR; bottom: western blot (5 pairs) **G**. Representative images of UCHL1 and p53 IHC staining in human serous tubal intraepithelial carcinoma (STIC); scale bar: 50µm and 20µm. **H**. Kaplan Meier survival curves for 1104 HGSOC patients with low or high UCHL1 levels after chemotherapy and optimal and suboptimal debulking. Progression-free survival was analyzed by KMplotter using auto-select best cut-off (p = 0.00021). **I**. Using OVMARK disease-free survival of HGSOC patients (n=244) with low or high UCHL1 levels was analyzed after optimal debulking and median expression cut-off in an *in-house* cohort of Molecular Therapeutics for Cancer, Ireland and GSE9899 (p = 0.004). Statistical significance was determined by the log-rank test, one-way ANOVA, and Student *t-*test. *p<0.05. The box boundaries represent the upper and lower quartiles, the horizontal line represents the median value, and the whiskers represent the minimum and maximum values.

### Epigenetic upregulation of UCHL1 promotes HGSOC growth

To understand the role of UCHL1 in HGSOC pathobiology, we examined the expression of UCHL1 in a panel of ovarian cancer cell lines [45] characterized as HGSOC and non-HGSOC cell lines. Compared to non-HGSOC, UCHL1 mRNA, and protein levels were significantly higher in HGSOC cell lines (Figure 2A). Interestingly, the elevated UCHL1 levels in HGSOC cells varied with the different *TP53* mutations and mutant p53 expression levels in these cell lines (Figures 2A and 2B). Similarly, a weak correlation (r = 0.2) was seen between UCHL1 and mutant p53 expression levels in HGSOC patients with missense p53 mutations (Figure S2A). In contrast, UCHL1 expression was low or absent in the non-HGSOC cells with WT p53 or p53-null respectively (Figures 2A and 2B). These results confirm our patient data and suggest that UCHL1 expression is not epigenetically silenced in HGSOC as reported in many malignancies [25, 26]. To test this, we performed methylated DNA immunoprecipitation (MeDIP) using 5-methylcytosine (5MC) antibody in Kuramochi and OVCAR3 (HGSOC) and HeyA8 and OVCAR5 (non-HGSOC) cells. No enrichment of methylated DNA in the UCHL1 promoter was observed in HGSOC cells, while significant enrichment was observed in non-HGSOC cell lines (Figure 2C and S2B). Chromatin immunoprecipitation (ChIP) assay using the H3K4 trimethylated antibody revealed enhanced enrichment of H3K4 trimethylated chromatin in the UCHL1 promoter in HGSOC cells, OVCAR3 and OVCAR4 (Figure 2D). However, no such enrichment of H3K4 trimethylated chromatin was observed in SKOV3 (Figure 2D). Furthermore, open chromatin marks at the UCHL1 gene promoter (chromosome 4; region 41257000-41258000; exon 1 to exon 3) were revealed by ATAC-seq analysis of OVCAR3 cells, unlike the non-HGSOC, SKOV3 cells (Figure 2E). To further corroborate these results, we next treated HGSOC and non-HGSOC cell lines with DNA methyltransferase inhibitor, 5-Aza-2’Deoxycytidine (5-Aza-DC). No change in UCHL1 expression was observed in HGSOC cell lines upon treatment with 5-Aza-DC (Figure S2C), while UCHL1 expression was increased many folds in non-HGSOC cell lines (Figure S2D). Similarly, 5-Aza-DC treatment in FT epithelial cells demonstrated a significant increase in the UCHL1 (Figure S2E). Collectively, these indicate the presence of unmethylated CpG islands in the UCHL1 promoter and epigenetic upregulation of UCHL1 in HGSOC.

**Figure 2.**
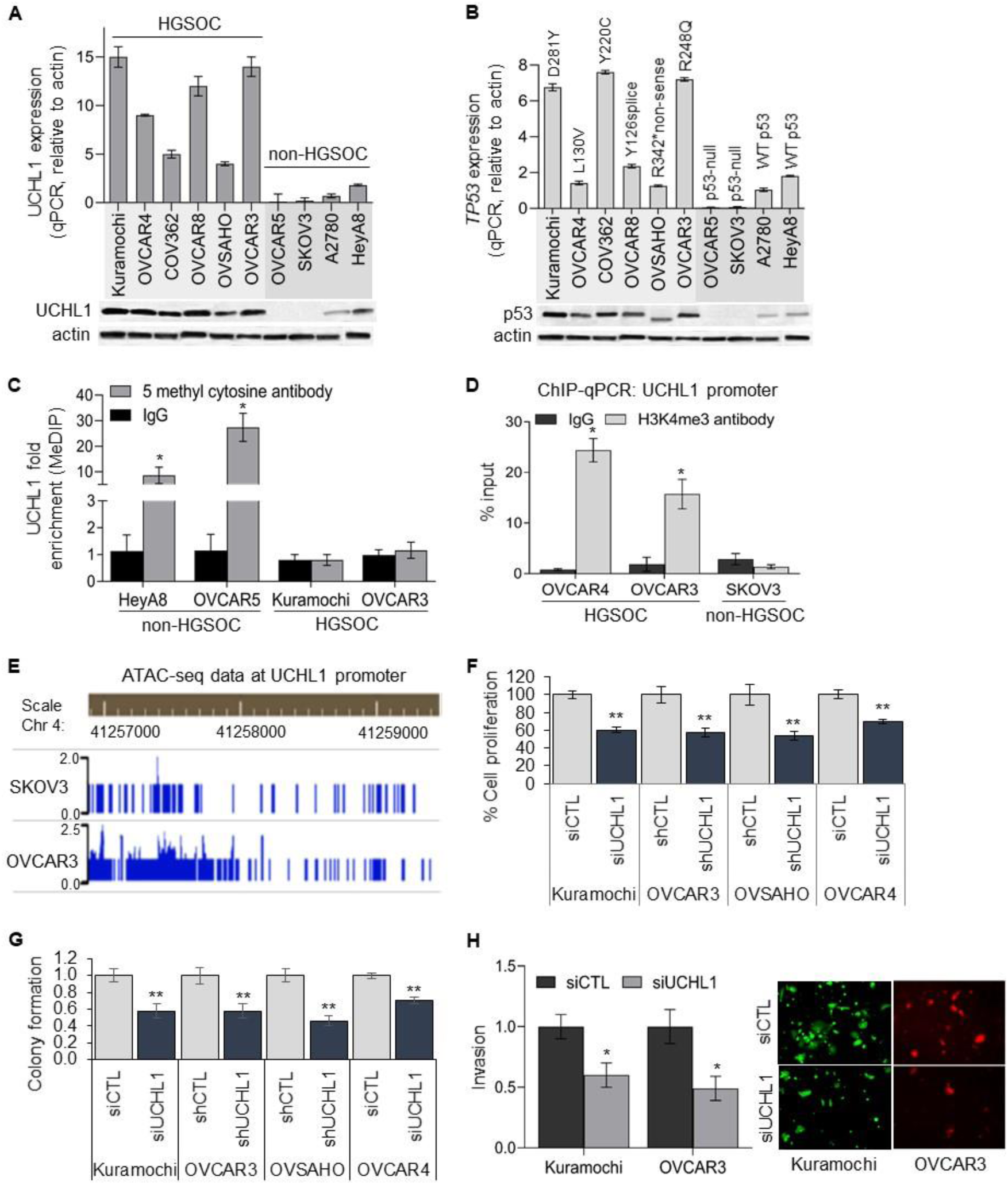
Epigenetic upregulation of UCHL1 promotes HGSOC growth. **A-B**. UCHL1 and p53 mRNA and protein levels in HGSOC and non-HGSOC cells. Respective p53 mutation status is given above the bars showing p53 mRNA expression in (B). **C**. Methylated DNA immunoprecipitation (MeDIP) was performed using 5 methylcytosine antibody or control IgG in HGSOC and non-HGSOC cells followed by qPCR for UCHL1 promoter. Methylated DNA enrichment in the UCHL1 promoter is shown relative to control IgG. **D**. Chromatin immunoprecipitation (ChIP) assay was performed using anti-histone H3 trimethyl lysine 4 (H3K4me3) antibody or control IgG in HGSOC and non-HGSOC cells followed by qPCR for UCHL1 promoter. H3K4 trimethylated chromatin enrichment in the UCHL1 promoter is shown relative to the input. **E**. ATAC-seq sequencing tracks at the UCHL1 gene locus in OVCAR3 and SKOV3 cells. Each track represents chromatin accessibility per 100bp bin. The region shown is human chromosome 4 (chr4):41257000-41259000. **F-G** Relative proliferation and clonogenic growth of HGSOC cells; Kuramochi, OVCAR3, OVCAR4, and OVSAHO transfected or transduced with control or UCHL1 siRNA and control or UCHL1 shRNA lentiviral particles. 2000 cells/well were plated in the 96 well plates and MTT assay was performed on day 4. 1000 cells/well were plated in the 6-well plates and colonies were fixed, stained by crystal violet after 8-10 days. **H**. Invasion of OVCAR3 (RFP labeled) and Kuramochi (GFP labeled) cells transfected with control or UCHL1 siRNA through the layers of normal human omental primary mesothelial cells and fibroblasts in a transwell insert (8µm pore size). Invaded cells were fixed after 16h, imaged and counted. Statistical significance was determined by Student *t-*test from at least three independent experiments. *p<0.05, **p<0.001. See Supplementary Figure S2.

To understand the functional effects of UCHL1 in HGSOC, we knocked down UCHL1 in HGSOC cell lines: Kuramochi, OVCAR3, OVCAR4, and OVSAHO (Figure S2F). Cellular proliferation (Figure 3A) and clonogenic growth (Figures 3B and S2G) of HGSOC cells were significantly reduced upon silencing UCHL1. Next, we studied the effect of UCHL1 silencing on the invasion of HGSOC cells. Omentum is the most favorable site for HGSOC metastatic growth. To mimic the invasion of cancer cells through the outer layers of the omentum during metastasis, we utilized an organotypic three-dimensional (3D) culture model of the omentum assembled in a transwell insert (Figure S2H) [35]. The invasion of UCHL1 silenced Kuramochi (GFP labeled) and OVCAR3 (RFP labeled) cells through the omental cells was significantly reduced compared to the unsilenced controls (Figure 3D). Together, this data demonstrates that UCHL1 promotes growth and invasion.

**Figure 3.**
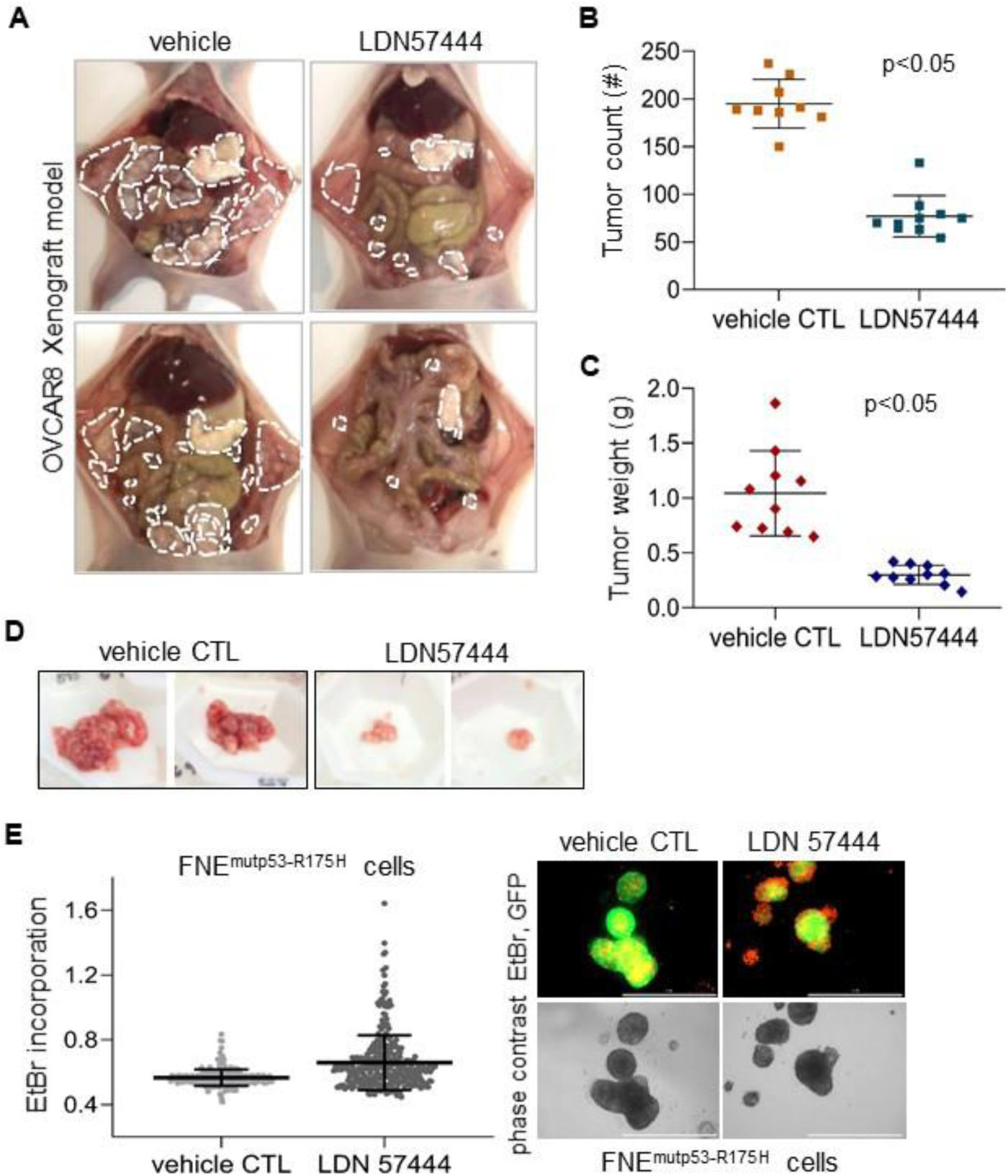
Effect of UCHL1 inhibitor, LDN57444 on HGSOC metastatic growth. **A**. Representative images of metastatic tumor colonies (encircled in dotted line) in the athymic nude mice received the intraperitoneal injection of OVCAR8 cells, and treated with vehicle control or UCHL1 inhibitor, LDN57444 (LDN) 1mg/Kg thrice per week (n=10 per group). **B**. Number of tumor nodules in mice treated with LDN or vehicle control. **C**. Weight of surgically resected tumors in the vehicle control and LDN-treated mice. **D**. Representative images of surgically resected tumors in the vehicle control and LDN-treated mice. **E**. Fallopian tube non-ciliated epithelial (FNE) cells transfected with pWZL-p53-R175H were cultured in ultra-low attachment plates and treated with vehicle control or UCHL1 inhibitor, LDN57444 (LDN) 10µM, 5 days. Ethidium bromide (EtBr) incorporation was quantified after 5 days in cellular clusters treated with DMSO (n=254) and LDN (n=334). Representative images of DMSO or LDN-treated cellular clusters of GFP labeled FNE^mutp53-R175H^ cells, scale bar 1000µM. EtBr incorporation is visible as orange color. Data are represented as mean ± standard deviation. Statistical significance was determined by Student *t-* test, *p<0.05.

### UCHL1 inhibitor, LDN57444 inhibits HGSOC metastatic growth

To investigate the effect of UCHL1 on tumor growth *in vivo*, we treated a mouse xenograft model of HGSOC metastasis with the UCHL1 inhibitor, LDN57444 (LDN). Athymic nude mice were intraperitoneally injected with 5 million OVCAR8 cells and peritoneal metastases were allowed to form. Subsequently, mice were intraperitoneally treated with LDN (1mg/Kg) or vehicle control thrice per week (10 mice/group). Mice were euthanized 45 days after injecting the cancer cells and the tumors were counted, surgically resected, and weighed. LDN treatment resulted in significantly smaller and fewer metastases compared to vehicle controls (Figures 3A and 3B). Furthermore, the overall weight of the surgically resected tumors was significantly less in the LDN-treated mice compared the control mice (Figures 3C and 3D). Hematoxylin and eosin staining of the tumor sections revealed that the tumors from the LDN and control groups were histologically similar (Figure S3A). These results demonstrate the potential of the UCHL1 inhibitor, LDN57444 in abrogating metastatic growth *in vivo*. Similarly, *in vitro* treatment with LDN as well as UCHL1 knockdown in OVCAR8 cells significantly reduced the cell growth (Figures S3B and S3C). On the contrary, LDN treatment in OVCAR5 cells (with no endogenous UCHL1 expression) showed no effect on cellular proliferation (Figure S3D) demonstrating the specificity of LDN57444 for UCHL1.

HGSOC precursor lesions in the fallopian tube (FT) uniquely disseminate through the peritoneal fluid, which largely depends on the anchorage-independent survival of cancer cells. Therefore, we next studied the effect of UCHL1 inhibitor, LDN on the anchorage-independent survival using a model of such early dissemination consisting of spheroids of FT non-ciliated epithelial (FNE) cells transfected with pWZL-p53-R175H (FNE^mutp53-R175H^). Compared to empty vector controls, prolonged anchorage-independent survival of FNE^mutp53-R175H^ spheroids, overexpressing mutant p53 variant R175, has been reported previously [34]. We observed increased expression of UCHL1 in FNE^mutp53-R175H^ cells (Figure S2E). The growth of FNE^mutp53-R175H^ spheroids was significantly reduced upon LDN (10µM, 5 days) treatment (Figure 3E) as evidenced by the increased ethidium bromide (EtBr) intercalation into DNA due to cell death associated with nuclear membrane fracture and reduced GFP expression in the LDN-treated FNE^mutp53-R175H^ spheroids compared to untreated controls (Figure 3E). These results indicate that UCHL1 inhibition increases flotation-induced cell death. Collectively, the data demonstrate that UCHL1 affects HGSOC metastatic growth.

### UCHL1 knockdown results in the activation of unfolded protein response and impair the proteasome activity

UCHL1 has been known to have varied functions including DNA binding, promoting translation initiation, influencing gene expression [46-48]. Therefore, to get a better overall understanding of its mechanism of action in HGSOC, we conducted RNA-seq analysis in the UCHL1 silenced Kurmamochi cells. A total of 1004 genes were significantly differentially expressed in Kuramochi cells upon silencing UCHL1 with a 1% false discovery rate. Analysis of the top 35 dysregulated genes (Figure 4A) revealed the upregulation of stress-induced genes, including Heme Oxygenase 1 (HMOX1) – a heat-shock factor 1 (HSF1) target gene [16] and activating transcription factor 3 (ATF3) – an endoplasmic reticulum (ER) stress-induced gene. Volcano plot of differentially expressed genes revealed activation of unfolded protein response (UPR) as measured by the upregulation of DDIT3 (CHOP), ATF4, ATF3, GADD35, HSP40 [16] (Figure 4B). In contrast to the upregulation of stress-induced genes, the genes implicated with proteasome activity were the top down-regulated genes in our RNA-seq data (Figure 4A). The expression of proteasome subunit alpha 7 (PSMA7) and acylaminoacyl peptide hydrolase (APEH) was significantly reduced upon silencing UCHL1 in our RNA-seq data (Figure 4A) and subsequent validation by qPCR in UCHL1 silenced HGSOC cell lines (Figure 4C). Inhibition of proteasome activity has been associated with the induction of terminal UPR [9, 49]. Our data indicate that UCHL1 inhibition results in activation of the ER stress or proteotoxic stress response potentially due to impaired proteasome activity and degradation of proteins.

**Figure 4.**
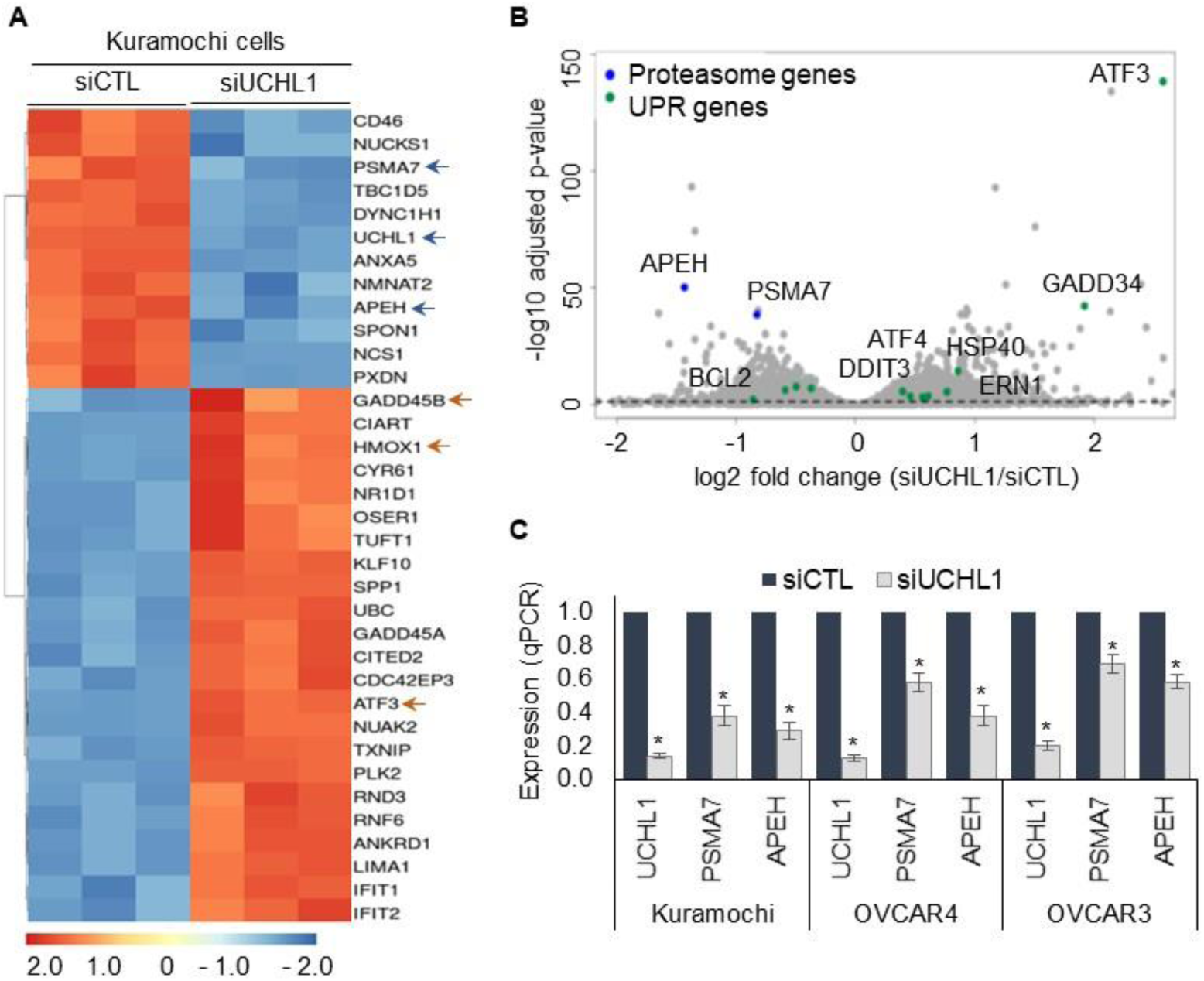
UCHL1 knockdown affects proteasome function and triggers the unfolded protein response. **A**. Heat map of the top 32 differentially expressed genes identified by RNA-sequencing of Kuramochi cells transfected with control or UCHL1 siRNA (p<0.005; 1% FDR). Orange color represents upregulated genes, while blue color represents downregulated genes. **B**. Volcano plot represents the global effect of UCHL1 knockdown on the expression of genes involved in the adaptive response. The x-axis shows the log2 fold change in the expression of genes between control siRNA and UCHL1 siRNA transfected Kuramochi cells; the y-axis shows the p-value for that difference. The dots on the negative and positive values of the x-axis represent downregulated and upregulated genes respectively. The dashed line represents p=0.05 at 1% FDR. Genes falling above the dashed line are significantly changed. Blue dots represent genes implicated with proteasome activity, while the green dots represent ER-stress induced genes. **C**. PSMA7 and APEH expression (qPCR) in Kuramochi, OVCAR4, and OVCAR3 cells transfected with control or UCHL1 siRNA.

### PSMA7 and APEH mediates proteasome activity and HGSOC growth

The proteasome subunit alpha 7 (PSMA7) proteasome isoform has been associated with enhanced resistance to stress in yeast and primed mammalian cells [50]. To evaluate the significance of PSMA7 in HGSOC, we analyzed PSMA7 expression in HGSOC patient tumors in TCGA database. PSMA7 was found to be overexpressed in HGSOC tumors (Figure 5A) and correlated with poor overall survival of HGSOC patients after optimal debulking (Figure 5B). Silencing PSMA7 demonstrated significantly reduced chymotrypsin-like proteasome activity and 20S proteasome levels in HGSOC cells (Figures 5C and 5D), leading to the accumulation of polyubiquitinated proteins (Figure 5E). Consistent with these findings, cellular proliferation and clonogenic growth of PSMA7 silenced HGSOC cells were significantly reduced (Figures 5F and 5G). These results suggest that PSMA7-mediated proteasome activity is required for HGSOC growth. Similarly, the activity of cytosolic enzyme, APEH has been associated with increased proteasome activity [37]. APEH catalyzes the removal of *N*-acetylated amino acid from the acetylated peptides leading to the release of free amino acids. The activity of APEH possibly disrupts the negative feedback inhibition of proteasomal activity caused by the accumulation of *N*-acetylated peptides after proteasomal degradation of proteins [37]. The expression of APEH was significantly high in HGSOC tumors compared to normal ovaries in TCGA database (Figure 5H). Moreover, APEH activity and expression were significantly reduced in UCHL1 or APEH silenced HGSOC cells (Figures 5I, 5J, and 5K), and the reduced APEH activity decreased chymotrypsin-like proteasome activity in HGSOC cells (Figure 5L). Supporting these results, cellular proliferation and clonogenic growth of HGSOC cells were significantly reduced upon silencing APEH (Figures 5M and 5N). Collectively, these results demonstrate that the UCHL1-PSMA7-APEH axis mediates proteasome activity and HGSOC growth.

**Figure 5.**
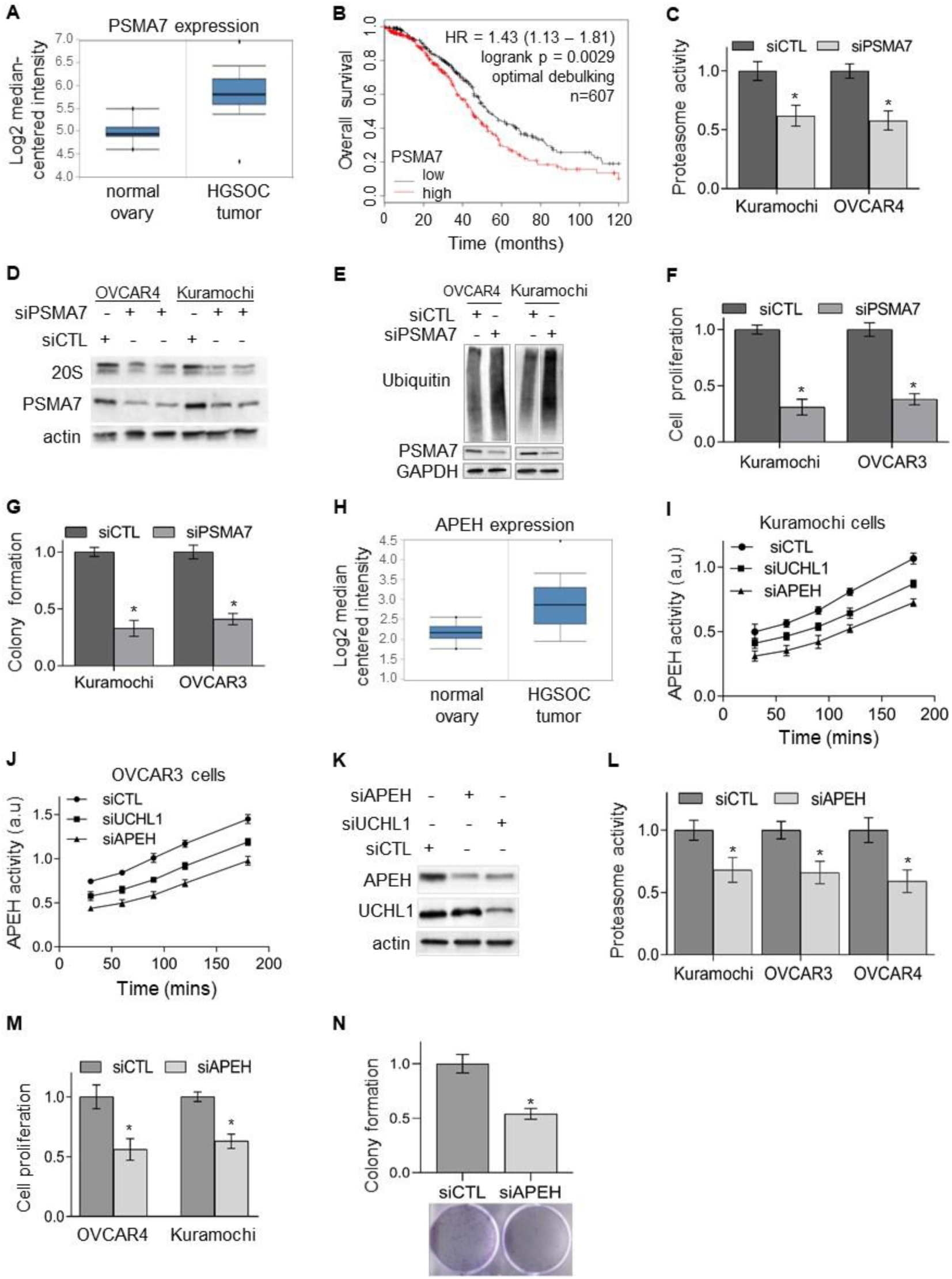
PSMA7 and APEH mediate proteasome activity and HGSOC growth. **A**. PSMA7 mRNA expression in normal ovary and HGSOC patient tumors in TCGA database analyzed using the Oncomine gene browser. **B**. Kaplan Meier survival curves showing overall survival of 607 HGSOC patients with low or high PSMA7 levels after optimal debulking and chemotherapy analyzed by KMplotter (p=0.0027). **C**. Chymotrypsin-like proteasome activity was measured using fluorescent substrate LLVY-R110 in cell lysates of OVCAR4 and Kuramochi cells transfected with control or PSMA7 siRNA. The cleavage of LLVY-R110 by proteasomes was monitored fluorometrically. **D-E**. Representative immunoblot analysis of 20S proteasome, PSMA7, and total ubiquitinated proteins in OVCAR4 and Kuramochi HGSOC cells transfected with control or PSMA7 siRNA. **F-G**. The relative proliferation and clonogenic growth of HGSOC cells transfected with control or PSMA7 siRNA. 2000 cells/well were plated in the 96 well plates and MTT assay was performed on day 4. 1000 cells/well were plated in the 6-well plates and colonies were fixed, stained by crystal violet after 8-10 days. **H**. APEH mRNA expression in normal ovary and HGSOC patient tumors in TCGA database analyzed using the Oncomine gene browser. **I-J**. APEH activity was measured by acetyl‐Ala‐*p*‐nitroanilide (Ac‐Ala‐pNA) in the total cell lysate of HGSOC cells transfected with control or UCHL1 or APEH siRNA. The cleavage of colorimetric p-nitroanilide by APEH was measured at different time points. **K**. Representative immunoblot analysis of APEH and UCHL1 in the whole cell lysate of Kuramochi cells transfected with control or UCHL1 or APEH siRNA. **L**. Chymotrypsin-like proteasome activity was measured using fluorescent substrate LLVY-R110 in OVCAR3, OVCAR4, and Kuramochi cells transfected with control or APEH siRNA. The cleavage of LLVY-R110 by proteasomes was monitored fluorometrically. **M-N**. The relative proliferation and clonogenic growth of Kuramochi and OVCAR4 cells transfected with control or APEH siRNA. 2000 cells/well were plated in the 96 well plates and MTT assay was performed on day 4. 1000 cells/well were plated in the 6-well plates and colonies were fixed, stained by crystal violet after 8-10 days. Statistical significance was determined by unpaired Student *t-*test from at least three independent experimental repeats, *p<0.05. The box boundaries represent the upper and lower quartiles, the horizontal line represents the median value, and the whiskers represent the minimum and maximum values.

### UCHL1 inhibition attenuates mTORC1 activity and induces a terminal stress response

Inhibition of proteasomal degradation of misfolded and damaged proteins results in proteotoxicity leading to activation of terminal UPR and attenuation of protein translation [9, 51, 52]. Therefore, we hypothesize that UCHL1 inhibition can potentially render HGSOC cells vulnerable through impaired proteasomal activity and generation of proteotoxicity. This hypothesis is supported by a dose-dependent decrease in the proliferation of Kuramochi cells upon inhibiting proteasome activity by the second-generation proteasome inhibitor, carfilzomib (Figure 6A), and reduced proteasome activity in UCHL1 inhibitor (LDN57444) treated xenograft tumors (Figure 6B). Furthermore, both UCHL1 silencing, and treatment with LDN57444 resulted in the accumulation of polyubiquitinated proteins in HGSOC cells, Kuramochi, and OVCAR4 (Figures 6C and 6D). Consistent with these results, UCHL1 inhibition resulted in reduced mTORC1 (mammalian target of rapamycin complex 1) activity and protein synthesis as evidenced by decreased phosphorylated levels of two mTORC1 substrates, ribosomal protein S6 (S6), and the eukaryotic initiation factor 4E-binding protein (4EBP1) in UCHL1 silenced Kuramochi cells (Figure 6E). In contrast, the expression of ER stress-induced proteins, ATF4, ATF3, and pro-apoptotic protein CHOP was increased while the expression of anti-apoptotic protein BCL2 was decreased (Figure 6E). These results indicate that UCHL1 inhibition results in impaired protein degradation, leading to the accumulation of proteins, attenuation of protein synthesis, and activation of terminal UPR. Collectively, the data demonstrate that UCHL1 promotes HGSOC growth by mediating proteasomal degradation of misfolded proteins through the PSMA7-APEH-proteasome axis and maintains protein homeostasis. Inhibiting UCHL1 results in proteotoxicity and activates terminal UPR (Figure 6F).

**Figure 6.**
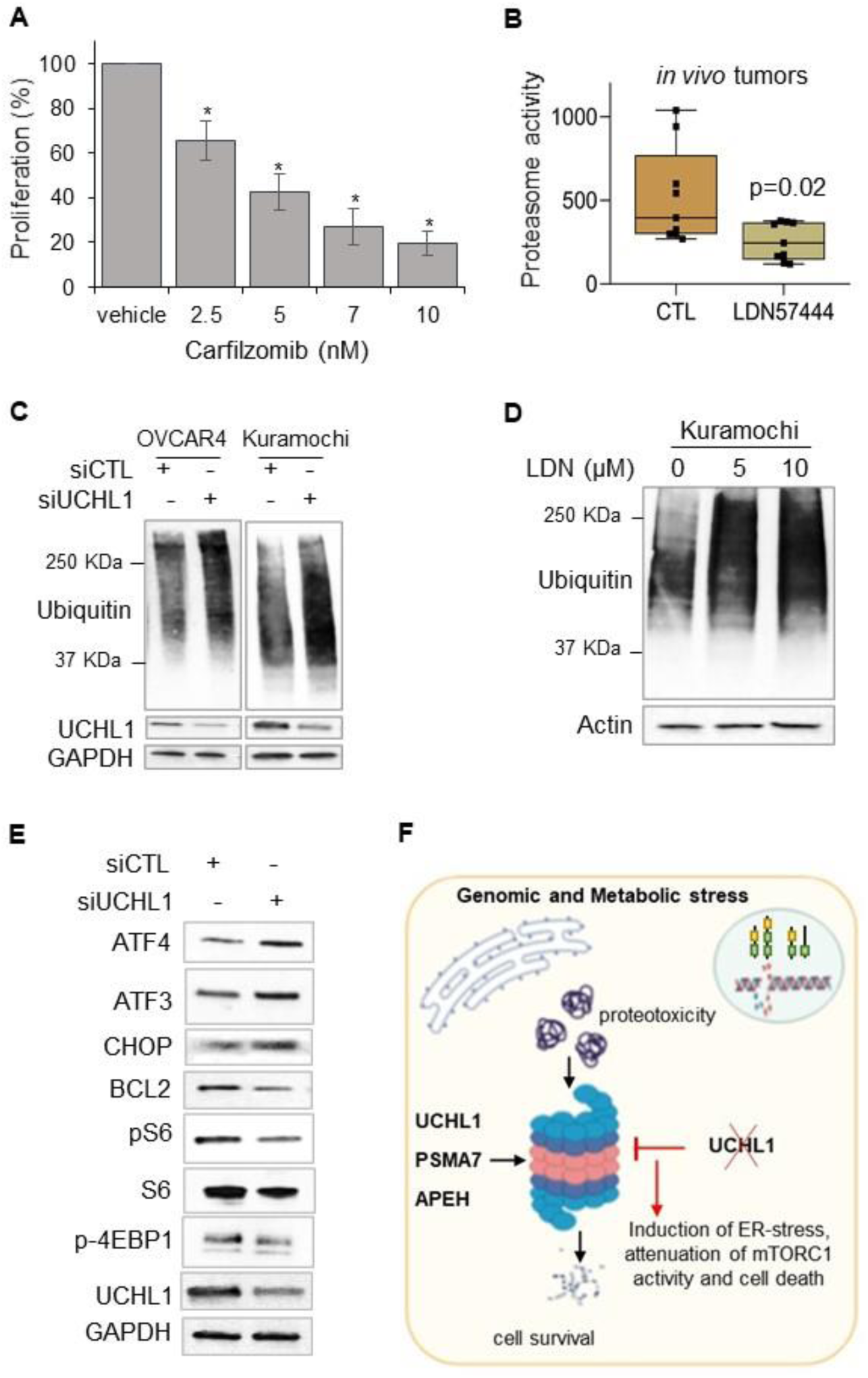
UCHL1 inhibition attenuates mTORC1 activity and induces a terminal ER stress response. **A**. Relative proliferation of Kuramochi cells treated with vehicle control or proteasome inhibitor, carfilzomib was measured by MTT assay after day 3. **B**. Chymotrypsin-like proteasome activity was measured using substrate LLVY-R110 in the tissue homogenate of xenograft tumors treated with the vehicle control or UCHL1 inhibitor, LDN 57444 (LDN). The cleavage of LLVY-R110 by proteasomes was monitored fluorometrically. **C**. Representative immunoblot analysis of total ubiquitinated proteins in OVCAR4 and Kuramochi HGSOC cells transfected with control or UCHL1 siRNA. **D**. Representative immunoblot analysis of total ubiquitinated proteins in Kuramochi cells treated with vehicle control or UCHL1 inhibitor, LDN57444 (5 and 10 µM) for 24 hrs. **E**. Representative immunoblot analysis of target proteins in Kuramochi cells transfected with control or UCHL1 siRNA. **F**. The schematic showing the role of the UCHL1-PSMA7-APEH-proteasome axis in mediating protein homeostasis and HGSOC cell survival. UCHL1 inhibition results in impaired proteasome activity and protein degradation leading to the accumulation of proteins, reduced mTORC1 activity and translation, and induction of UPR-associated apoptosis.

## Discussion

DUBs have been implicated in the regulation of many processes associated with tumor progression and are emerging as prognostic markers due to their correlation with tumor grade and stage [21, 53, 54]. UCHL1 is a cancer-associated DUB, which has been reported as either an overexpressed oncogene [21-23, 55] or epigenetically silenced tumor suppressor [25, 26, 56] in several malignancies. Previous studies have reported the role of UCHL1 in promoting metastasis by its deubiquitinating activity associated with HIF1α, cyclin B1, and TGFβ receptor 1 [21, 22, 24]. In this study, we have demonstrated that UCHL1 overexpression in HGSOC patients predicts poor prognosis and it promotes tumor growth by mediating protein homeostasis through the PSMA7-APEH-proteasome axis. Furthermore, we showed that inhibiting UCHL1 increases ER stress and induces terminal UPR due to impaired proteasome activity and accumulation of polyubiquitinated proteins. Previous studies have reported the induction of proteotoxic stress and cancer cell death by broadly inhibiting DUBs using a pan-deubiquitinating enzyme inhibitor and inhibitor of proteasome-associated DUBs [16, 57]. We have identified a specific DUB that mediates protein homeostasis, potentially through its association with proteasome, or cooperation with the UPR mediated pro-survival signaling. Moreover, about 96% of HGSOC patients harbor *TP53* mutations. Previous studies have reported the role of the mutant p53-NRF2 axis in transcriptional upregulation of proteasomal machinery [36] and the mutant p53-HSF1 (heat shock factor 1) axis renders cancer cells more resistant to proteotoxic and ER stress [58, 59]. Our patient data showed a weak correlation (r=0.2) between UCHL1 and mutant p53 expression levels in HGSOC patients with missense *TP53* mutations. Increased UCHL1 expression was also observed in STICs, and HGSOC cell lines harboring *TP53* mutations. Together, these findings indicate the context-dependent upregulation of UCHL1 in HGSOC. HGSOC originates from the FT secretory epithelial cells (FTSEC) [51]. The presence of abundant rough ER and well-developed Golgi complexes with secretory vesicles in FTSEC, a feature that remained in the malignant state [51], indicates that these cancer cells are primed for high protein synthesis, which renders them dependent on protein quality control pathways [51]. Moreover, profound genomic complexities and physiological stressors affect the protein folding capacity resulting in ER stress. From a translational perspective, this indicates HGSOC vulnerability to the imbalances in protein homeostasis and our study identifies novel links in this proteostasis network.

UCHL1 is mainly a neuronal DUB, it constitutes about 1-2% of total brain proteins. The loss of UCHL1 has been implicated in neurodegenerative diseases resulting in the accumulation of neuronal protein aggregates due to impaired proteasomal degradation [19, 20]. However, the exact mechanism remains elusive. For the first time, we report that increased expression of PSMA7 and APEH in HGSOC regulates proteasome activity and their association with the UCHL1 mediated proteostasis. Upregulation of proteasome subunits (PSMA3, PSMB5, and PSMA7) or proteasome assembly factors promote resistance to the proteasome inhibitors and ER stress [12, 50, 60, 61]. Specifically, the evolutionarily conserved PSMA7 proteasome isoform has been shown to provide tolerance to metallic stress in yeast and oxidative stress in the mammalian cells primed for PSMA7 proteasome formation [50]. Similarly, APEH regulates proteasome activity by catalyzing the removal of N-acetylated amino acid from the acetylated peptides and possibly disrupting the negative feedback inhibition of proteasomal activity caused by the accumulation peptides [37]. Taking this alternative approach of determining the genes and pathways transcriptionally deregulated by UCHL1 silencing revealed the role of PSMA7 and APEH in regulating proteasomal activity and degradation. Further studies are needed to identify the mechanism of their transcriptional regulation and the role of UCHL1 in this process. Since UCHL1 has been reported to be involved in varied functions from its interaction with DNA to affecting gene transcription and translation initiation [46-48]. It could be involved in a direct mechanism regulating the transcription of these genes or via its effect on the proteasomal machinery and protein turnover. Ataxin 3 is an example of DUB, which acts as a transcriptional co-repressor and is involved in protein homeostasis [62].

UCHL1 has been reported as an epigenetically silenced tumor suppressor in several malignancies. A previous study [26] has reported UCHL1 as an epigenetically silenced gene in ovarian cancer, however, UCHL1 was methylated in only 1 out of 17 tumors they studied [26]. Furthermore, the information on the ovarian cancer histotype was not provided [26]. Our findings revealed consistent upregulation of UCHL1 in multiple HGSOC datasets, including TCGA. Furthermore, the analysis of TCGA methylation data showed hypomethylation at UCHL1 gene loci in serous ovarian cancer specimens (data not shown), which corroborates with our MeDIP, ChIP, and ATAC-seq data in HGSOC cell lines. These findings revealed hypomethylation at the UCHL1 promoter and its epigenetic upregulation in HGSOC, potentially due to open chromatin at the gene loci. Mutant p53 has been reported to upregulate the expression of H3K4 histone methyltransferases in breast cancer [33], which in turn governs open chromatin and hypomethylation at gene loci. This further suggests the HGSOC-specific upregulation of UCHL1.

Targeting protein homeostasis by directly inhibiting proteasome activity has been clinically successful in certain tumor types such as multiple myeloma, possibly owing to its dependence on protein quality control pathways due to the inherently high protein synthesis rate [7]. Furthermore, in solid tumors, such as lung, pancreas, and head and neck cancer, the second-generation proteasome inhibitor, carfilzomib has started to show better results due to greater selectivity and inhibitory potency for proteasome subunits, and an improved clinical safety profile than Bortezomib [7]. These reports suggest that targeting proteasome or DUBs to induce proteotoxic stress is a viable approach to treat cancers [16, 57]. Various small-molecule DUB inhibitors are emerging as a therapeutic modality for cancer treatment, such as USP14 inhibitor, VLX1570 in myeloma, NCT02372240 [63]. In the present study, we identified the role of DUB UCHL1 in mediating protein homeostasis in HGSOC and the potential of UCHL1 and APEH inhibitors in sensitizing cancer cells to the proteotoxic stress.

## Acknowledgments

This research was funded by Ovarian Cancer Research Alliance, grant# 544389 to Sumegha Mitra. This work was also supported the other two pilot grants from Ralph W. and Young Grace M. Showalter Trust and Biomedical Research Grant from Indiana University School of Medicine to SM. The authors are thankful to Dr. Ronny Drapkin, Perelman School of Medicine, University of Pennsylvania for fallopian tube epithelial cells and STICs. We thank IU Health Pathology Laboratory, Dr. George Sandusky, Pathology and Laboratory Medicine, Ms. Constance J Temm, Research Immunohistochemistry Facility for their help in the tissue microarray analysis, and immunohistochemistry. We thank the Center for Medical Genomics - Dr. Yunlong Liu, Yue Wang, and Xiaona Chu for ATAC-sequencing.

## Authors’ Contribution

Kinzie Lighty: Investigation and validation. Apoorva Tangri: Investigation, methodology, and validation. Jagadish Loganathan: Investigation and validation. Fahmi Mesmar: Methodology and validation. Ram Podicheti: Formal analysis and investigation. Chi Zhang: Formal analysis and investigation. Marcin Iwanicki: Methodology, investigation, and writing review and editing. Harikrishna Nakshatri: Supervision, methodology, and writing review and editing. Sumegha Mitra: Conceptualization, funding acquisition, resources, methodology, investigation, supervision, project administration, and writing review and editing.

---

**Supplementary Figure S1.**
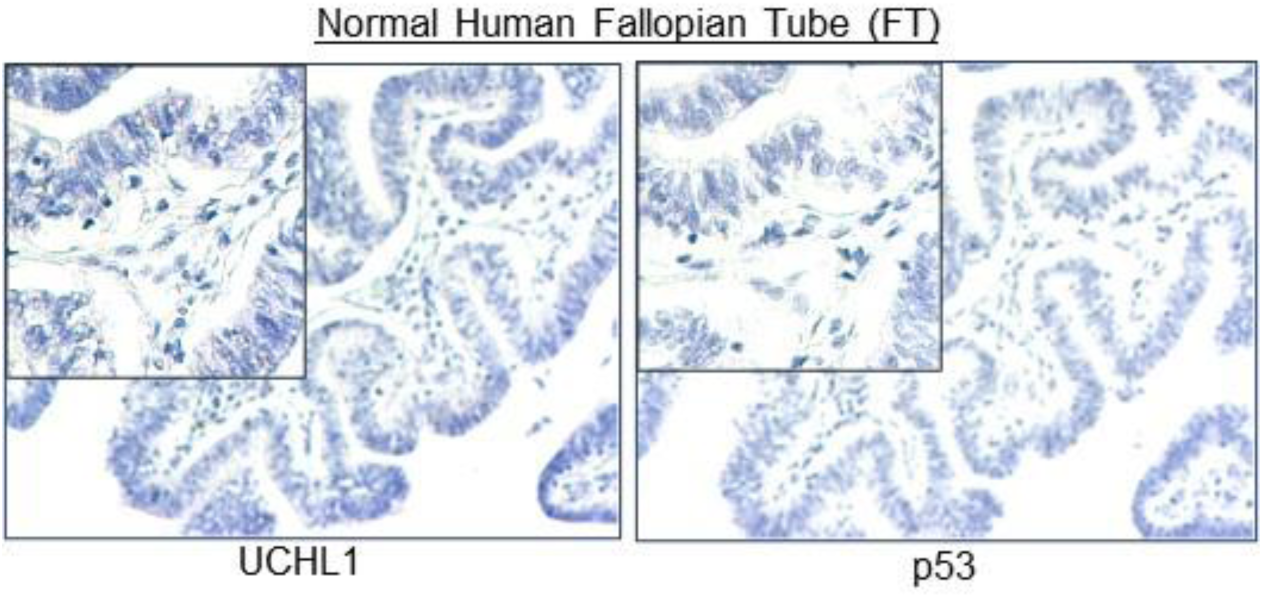
Representative image of UCHL1 and p53 immunohistochemical (IHC) staining in the normal human fallopian tube (n=5; 20x; scale bar: 50 µm and 150x magnified inset).

**Supplementary Figure S2.**
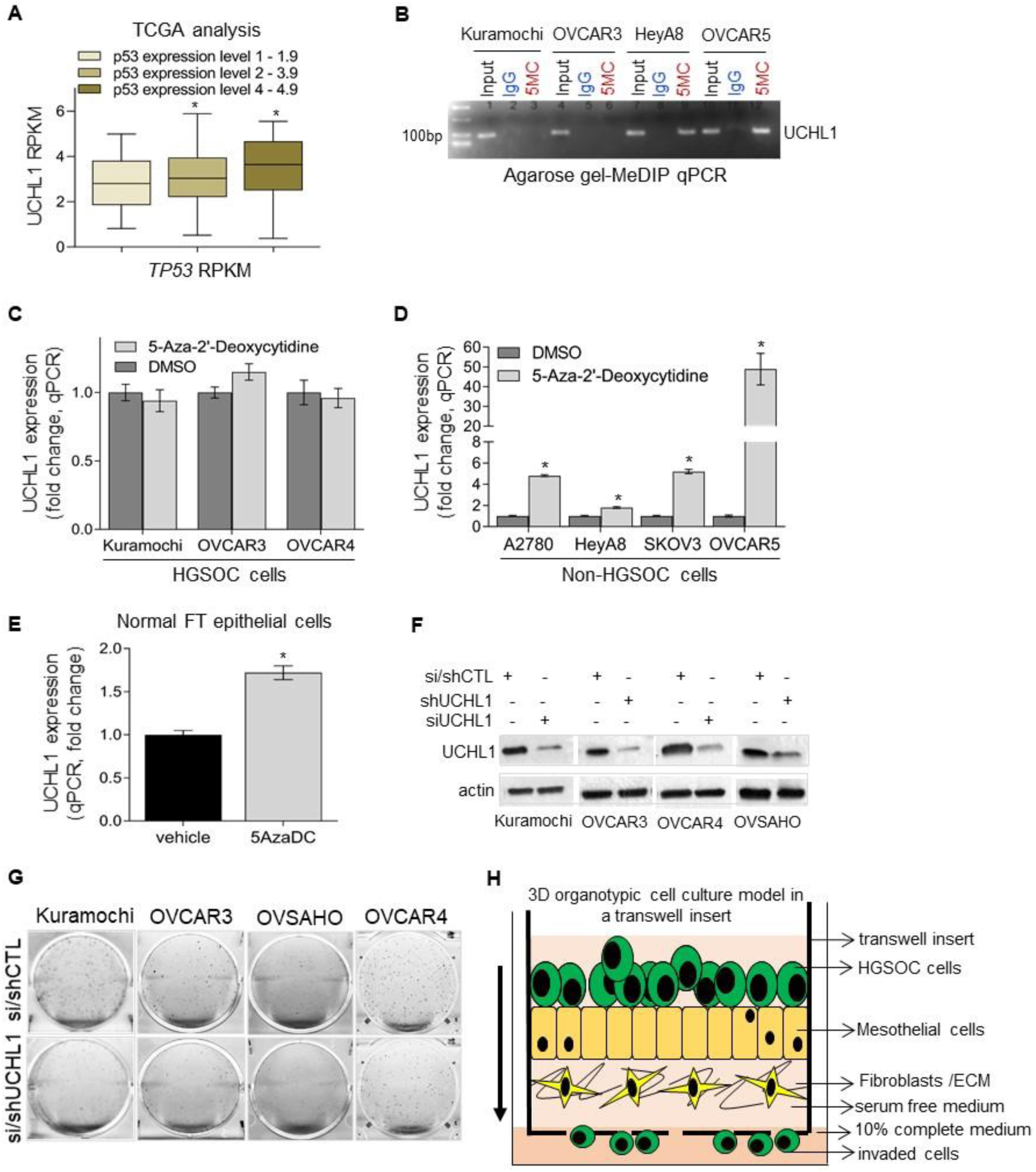
**A**. UCHL1 expression in HGSOC patients with missense *TP53* mutations and different p53 expression levels in the TCGA database analyzed using cBioportal. **B**. MeDIP-qPCR products were resolved using a 2% agarose gel with a 100bp ladder. The presence of amplicon with the 5MC antibody in non-HGSOC cell lines represents the enrichment of methylated DNA in the UCHL1 promoter. **C-D** UCHL1 expression (qPCR) in HGSOC and non-HGSOC cells treated with DNA methyltransferase inhibitor, 5-Aza-2’-deoxycytidine (5µM, 48h) or vehicle control. **E**. UCHL1 expression (qPCR) in normal human fallopian tube epithelial (FTE) cells treated with 5-Aza-2’-deoxycytidine (5µM, 48h) or vehicle control. **F**. Immunoblot analysis for UCHL1 in HGSOC cells transfected or transduced with control or UCHL1 siRNA and control or UCHL1 shRNA lentiviral particles. **G**. Representative images of colony formation assay of HGSOC cells transfected or transduced with control or UCHL1 siRNA and control or UCHL1 shRNA lentiviral particles. **H**. Schematic of invasion of HGSOC cells through 3D organotypic cell culture of omental primary human mesothelial cells and fibroblasts in a transwell insert. The insert was placed in a 24 well plate containing 10% complete medium and cancer cells invaded through omental cells towards the complete medium. Statistical significance was determined by Student *t-*test from at least three independent experiments. *p<0.05. The box boundaries represent the upper and lower quartiles, the horizontal line represents the median value, and the whiskers represent the minimum and maximum values.

**Supplementary Figure S3.**
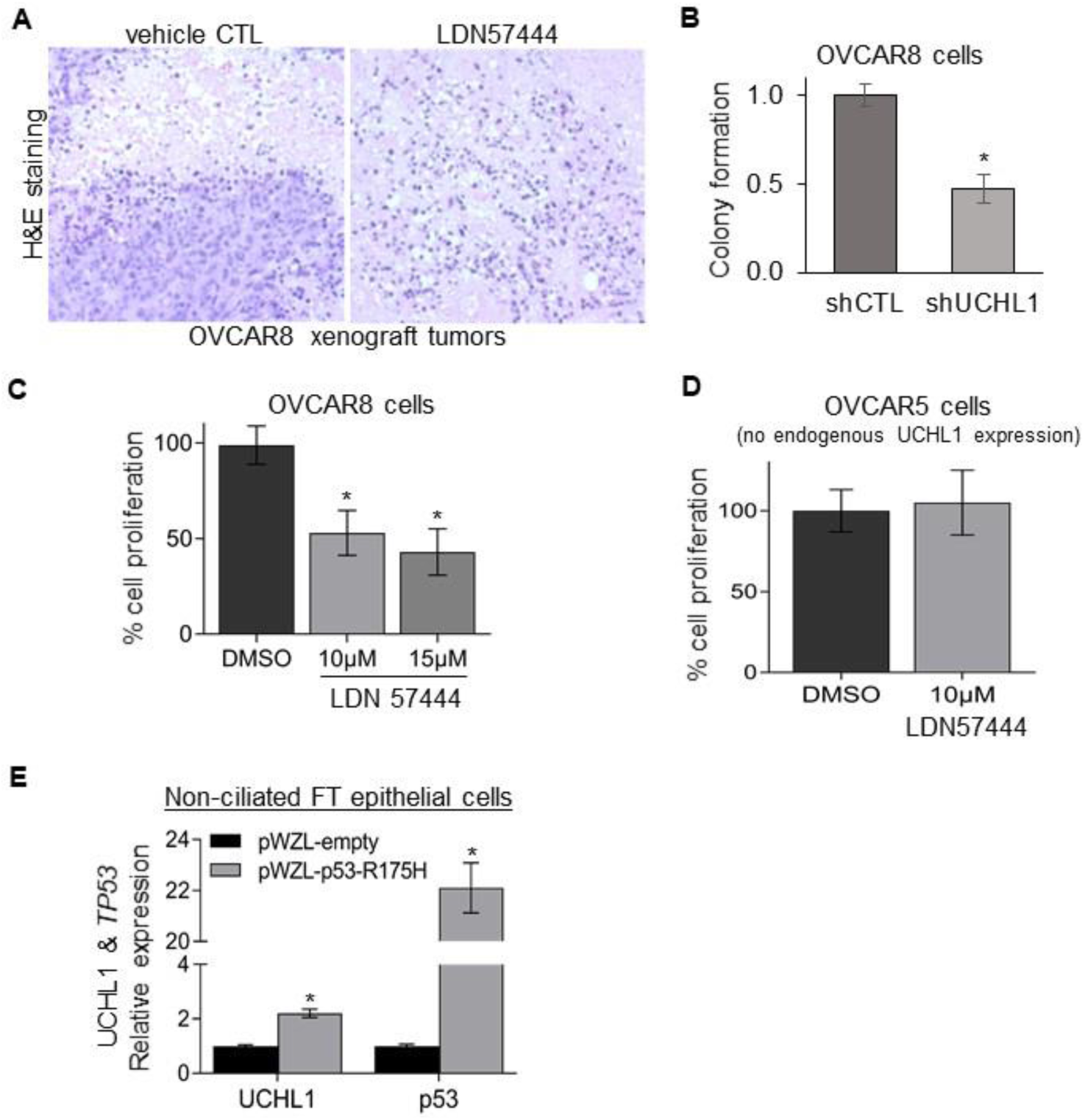
**A**. Representative images of hematoxylin and eosin staining of vehicle control and LDN-treated xenograft tumors. **B**. Relative clonogenic growth of OVCAR8 cells transduced with control or UCHL1 shRNA lentiviral particles. 1000 cells/well were plated in 6-well plates and the colonies were fixed, stained by crystal violet after 8 days. **C**. Relative proliferation of OVCAR8 cells treated with vehicle control or UCHL1 inhibitor, LDN57444 (LDN). 2000 cells were plated in the 96-well plates and treated with vehicle control and LDN (10µM) on day 1 and day 3, the MTT assay was performed on day 4. **D**. Relative proliferation of OVCAR5 cells (with nill UCHL1 expression) treated with vehicle control and LDN. 2000 cells were plated in the 96-well plates and treated with vehicle control and LDN (10µM) on day 1 and day 3, the MTT assay was performed on day 4. **E**. UCHL1 expression (qPCR) in the fallopian tube non-ciliated epithelial (FNE) cells transfected with empty vector or pWZL-mp53^R175H^. Data from at least 3 biological repeats, statistical significance was determined by Student *t-*test, *p<0.05.

